# Variational algorithms for analyzing noisy multi-state diffusion trajectories

**DOI:** 10.1101/278978

**Authors:** Martin Lindén, Johan Elf

**Affiliations:** Department of Cell and Molecular Biology, Uppsala University, Sweden.

## Abstract

Single particle tracking offers a non-invasive high-resolution probe of biomolecular reactions inside living cells. However, efficient data analysis methods that correctly account for various noise soures are needed to realize the full quantitative potential of the method. We report new algorithms for hidden Markov based analysis of single particle tracking data, which incorporate most sources of experimental noise, including heterogeneuous localization errors and missing positions. Compared to previous implementations, the algorithms offer significant speed-ups, support for a wider range of inference methods, and a simple user interface. This will enable more advanced and exploratory quantitative analysis of single particle tracking data.

---

Experimental techniques to track the conformational and binding states of single biomolecules can offer unique mechanistic insights into life at the molecular level, but increasingly rely on statistical computing to extract quantitative and reproducible results. A simple example is superresolved single particle tracking (SPT) [1], where changes in diffusion constant, or between different modes of motion, offers a non-invasive probe of binding and unbinding reactions in living cells [2, 3].

Accurate quantitative analysis of this type of data requires a faithful account of localization noise, which come in the form of localization errors and motion blur, sometimes referred to as “static” and “dynamic” errors, respectively [4, 5]. In particular, live cell imaging often lead to heterogeneous and asymmetric localization errors, for example due to photobleaching, variability between and. across cells, out-of-focus motion, or the dependence of localization errors on the diffusion constant [6, 7]. Several emerging techniques for 3D localization also give different precision in the axial and lateral directions [8].

A fundamental unknown in many live cell SPT studies is the number of underlying molecular states, which may differ in, e.g., diffusion constant. Counting diffusive states in SPT data presents a statistical model selection problem that has so far only been solved with simplified noise models [2], which may be inappropriate in many live cell applications [7, 9].

Here, we extend our previous HMM analysis [7] by deriving and implementing variational algorithms that increase computational speed by more than an order of magnitude, allow statistical model selection using Bayesian or information-theoretic methods, and can be generalized to a wider class of localization error models. The new methods are available in a user-friendly open source software suite.

## I. RESULTS

### A. Diffusive hidden Markov models with motion blur and localization uncertainty

The starting point for our analysis is a standard model for camera-based single particle tracking that includes a combination of averaging (motion blur) and localization errors, where the detected positions *x*_*t*_ are related to the underlying particle trajectory *y*(*t*) through

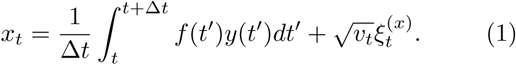

Here, *v*_*t*_ is the localization error (variance) in frame *t,* and 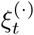 are independent *N* (0, 1)-variables. The shutter function *f* (*t*) describes how the image acquisition is distributed throughout the frame, e.g., *f* (*t*) = 1/Δ*t* for continuous acquisition [5].

We model the particle motion *y*(*t*) as free diffusion, with a time-dependent diffusion constant governed by a hidden Markov process *s*_*t*_ with *N* discrete states. With variational approximations in mind, we need a model in the exponential family to get an analytically tractable algorithms [10]. This means that we model both the true hidden path and the true exposure averaged positions (the integral in Eq. (1)) explicitly. In discrete time, this leads to the model

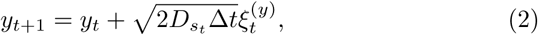

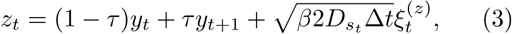

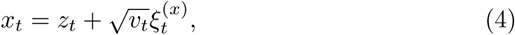

where *τ* and *β* are blur coefficients that depend on the shutter function [7] (see SI Sec. S1). Position coordinates in two- or three-dimensional trajectories are treated independently, which means that we neglect possible correlations between localization errors in different directions. As detailed below, and in the supplementary material, this model allows variational algorithms for both maximum likelihood estimation (MLE) and variational Bayes inference (VB) [11, 12].

Out focus in this work is the case where the localization variances *v*_*t*_ are input data, estimated from the localization of single spots [7]. However, one could also treat *v*_*t*_ as model parameters, for example as a single average error (*v*_*t*_ = *v*), dependent on the hidden state 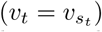, and/or varying with coordinate dimension. These modified models remain in the exponential family, and thus allow similarly efficient variational algorithms that differ only in details compared to our main case.

### B. Model selection

The number of diffusive states is often a biological unknown of great interest, but since different number of diffusive states correspond to statistical models with different number of parameters, counting states is a nontrivial problem of statistical model selection.

Bayesian reasoning, including model selection, is an extension of formal logic to uncertain statements, that yields unique and consistent results [13]. Assuming equal prior preference for a set of candidate models with uncertain parameters and unobserved degrees of freedom (latent variables), the Bayesian approach uses marginalization to select the model with the largest (log) evidence [11, 12], which in compact notation can be written

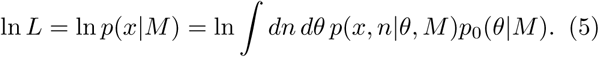

Here, *x* denotes the observed data, *M* the model, *n* = {*s*_*t*_, *y*_*t*_, *z*_*t*_} the latent variables (summed or integrated out as appropriate), *θ*_*M*_ the unknown parameters with prior distribution *p*_0_ (·|*M*) for the different models *M*.

In our case (and many others), the evidence is analytically intractable, in which case a variational approximation (“variational Bayes”, VB) can be an attractive approximative approach [2, 14–17]. VB yields an approximate log evidence usable for Bayesian model selection as well as approximate posterior distributions of parameters and hidden states [10–12]. Moreover, since it involves direct optimization of the evidence in Eq. (5), VB algorithms have an intrinsic parsimony that depopulates superfluous states and can be utilized for efficient greedy model search algorithms [2, 17].

However, Bayesian inference may be statistically inefficient. In particular, the common practice of using uninformative priors to minimize bias in parameter estimates means that the prior likelihood for any particular parameter value is low. This in turn can lead to overly steep penalties against models with many parameters, a phenomenon known as Lindley’s paradox, which means that an unnecessarily large amount of data is needed in order to resolve some feature of interest [18–20]. An alternative approach is to rank competing models by their estimated predictive performance, which does not suffer from Lindley’s paradox [19, 20]. The most famous example of this is Akaike’s information criterion (AIC) [21], but this is only asymptotically valid for regular models and large data sets. For finite data sets, one could insted use cross-validation, where the data is divided into two parts, one for learning model parameters (“training”) and one for estimating predictive performance (“validation”). In practice, the performance is estimated from averaging over several such divisions. Here, we focus on a Bayesian variant, pseudo-Bayes factors (PBF) [20], which are easy to compute with variational methods (see SI S2.5).

Figure 1 compares VB and PBF model selection on synthetic test data with three diffusive states and parameters that resemble in vivo SPT experiments in bacteria [2, 22] (see Methods). Broadly, one expects predictive model selection to avoid Lindley’s paradox and penalize complex models less severely than Bayesian methods as the amount of data increases, possibly at the expense of consistency, i.e., there is no guaranteed convergence to the correct model [23]. These expectations are qualitatively borne out in Fig. 1, where the Bayesian VB criterion is more prone to select too few states for small data sets, but less prone to select too many states for large data sets.

**Figure 1.**
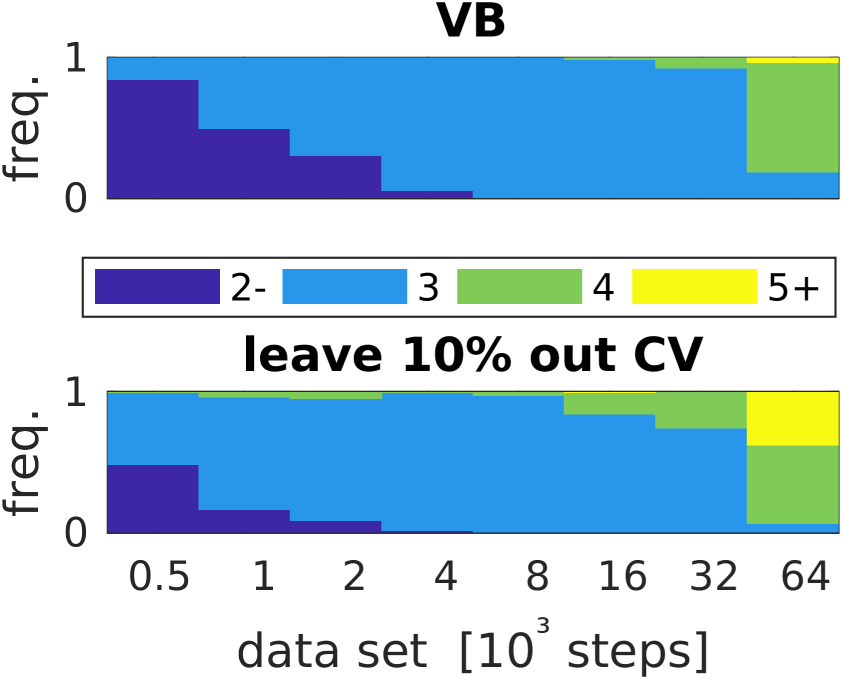
Statistical model selection. We generated a range of synthetic data sets with three diffusive states and *z*-dependent localization errors, and estimated the number of states using variational maximum evidence (top) and crossvalidation using variational pseudo-Bayes factors.

There is no simple optimal rule for selecting training and validation subsets, and HMMs also suffer the additional complication that individual observations are correlated due to the hidden state dynamics. SPT experiments typically generate many short trajectories, which we use as atoms for constructing training and validation sets. Fig. 1b uses randomly sampled validation sets containing about 10% of the data. Some other choices, including AIC, are explored in SI Fig. Figure S1, but turn out to perform worse.

### C. Speed-up

In addition to more flexibility in modeling and inference methods, the new algorithms are also considerably faster compared to our previous implementation [7]. This is mainly because variational algorithms based on Eqs. (2-4) are analytically tractable, and hence avoid a costly numerical optimization step, but also due to an improved matrix inversion algorithms [24]. Figure 2 shows the time per iteration for a 3-state model on data sets of different sizes for the new algorithm compared to that of Ref. [7]. We see speedups of 1-2 orders of magnitude for experimentally relevant data set sizes of 10^4^ 10^5^positions, as well as better scaling.

**Figure 2.**
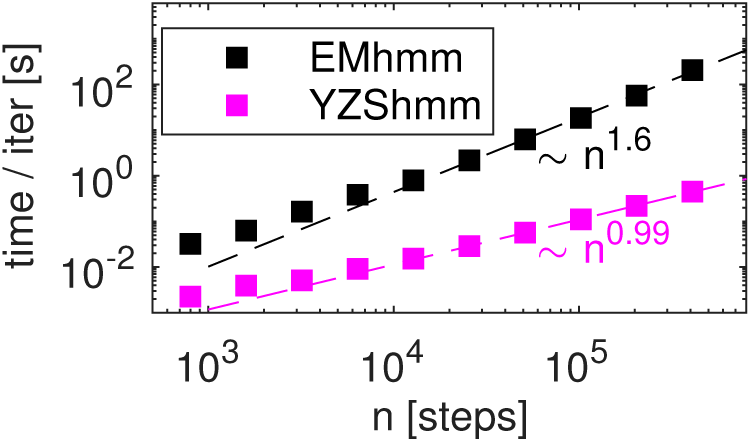
Speed of our variational algorithm (YZShmm) compared to that of Ref. [7] (EMhmm). Time per iteration vs. number of steps in the data (a trajectory with *T* positions contains *T -* 1 steps), with 3 diffusive states, was measured on a dual 6-core Intel Xeon 2.4GHz computer running Matlab R2017a. Scaling laws are guides to the eye.

### D. Finding the global optimum

Variational learning of a model and its parameters, diffusion constants and transition rates, involves finding the overall best fit to the data, but VB and other expectation-maximization type algorithms only converge towards local optima. An additional global search is needed.

The simplest approach is to converge multiple models from different starting points. To speed things up, we use the built-in parsimony of the VB algorithms to start from complex many state models and then systematically search for simpler ones by removing un or lowpopulated states [2]. However, the extra complexity of our model compared to standard HMMs [2] makes this approach more challenging to apply.

One attractive feature of our model is the ability to handle long trajectories with missing positions due to, e.g., fluorophore blinking, by setting *v*_*t*_ = *∞*. However, when fitting high-dimensional models to data with missing positions, groups of superfluous states sometimes converge towards identical parameters and finite occupancy associated with the missing positions. Since this is clearly unphysical, we choose to remove such state clusters before commencing normal model pruning.

Another challenge is related to the presence of two types of latent variables, for the discrete diffusive states (*s*_*t*_) and uncertainty in true particle positions (*y*_*t*_, *z*_*t*_), respectively. Although hard to quantify, it seems reasonable to expect more latent variables to yield a more complex search landscape, with more local optima for the search to get trapped in, compared to ordinary HMMs where the particle positions are not latent variables [2]. More concretely, the variational treatment uses a three-fold factorization ansatz (parameters, hidden states, and hidden particle trajectories), and to initialize the local optimization iterations, two of three factors need to be initialized.

We use randomly selected parameter values, and explore different strategies to initialize either hidden states or hidden trajectories: uniform hidden state occupancy, hidden trajectories modeled directly on observed data (with no uncertainty or correlations between *y*_*t*_ and *z*_*t*_), and hidden trajectory models generated by a running average where a pure diffusion model is fit to a small window of variable length. Generally, there does not seem to be a single silver bullet. Instead, differnt initialization methods do well on different types of data. Figure 3 shows an analysis of a data set from simulated images (see Methods) using 50 independent initializations of model parameters with 15 states, and 10 different initializations of latent variables with each parameter set. Fig. 3a shows the lower bounds of models originating from a single initial parameter set, with each line corresponding to models generated by the reductive search starting from one latent variable initialization. We see that the initialization with the largest number of non-spurious states does not lead to the best overall model, and that the search lines sometimes cross, meaning that relative ranking among the different reduction searches can change as states are removed. Looking at the best models from 50 independent parameter initializations (Fig. 3b), we again see search lines crossing, and note that the two most high ranked model sizes originate from different initialization methods.

**Figure 3.**
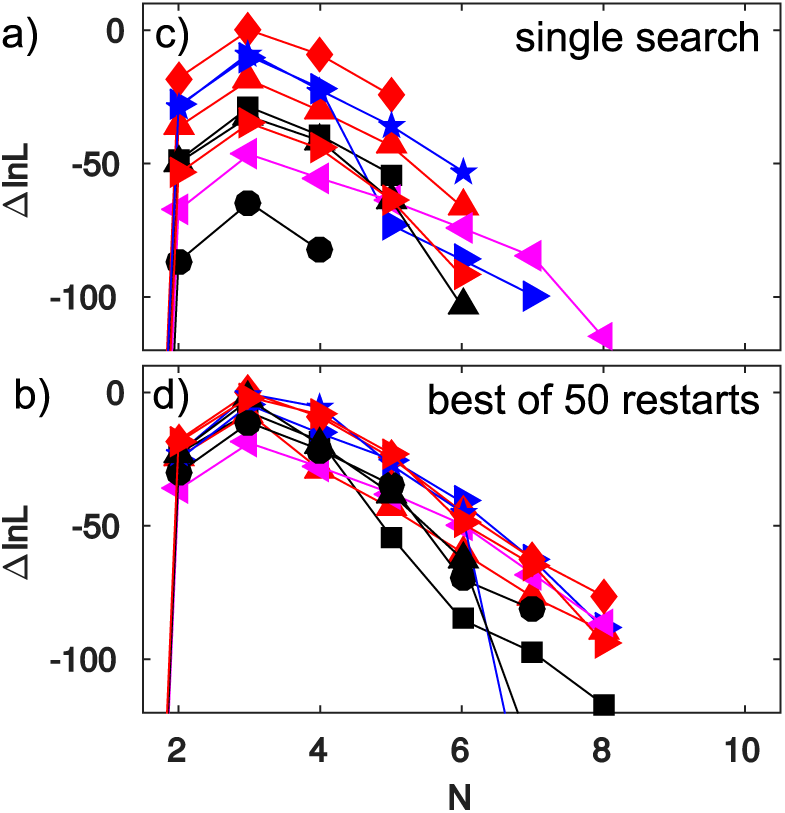
Model search with different initialization strategies. Each color/marker combination shows to the lower bounds *lnL* from the best model of each size found from different latent variable (hidden states *s*_*t*_ or position uncertainties *y*_*t*_, *z*_*t*_) initializations. (a) Model search from a single parameter initialization. (b) Best models from 50 independent restarts.

### E. Application: simulated tRNA tracking

Quantitative live cell single particle tracking is complex, and errors may arise during measurement, spot detection, localization, trajectory building, and trajectory analysis. Comprehensive tests of the whole analysis chain are needed to validate quantitative interpretations of the experiments under particular conditions. To evaluate the capabilities of our new trajectory analysis only, we seek test data with known ground truth and sufficient realism to be experimentally relevant. We use simulated video microscopy [9] to produce realistic test data, run spot detection and localization using our standard methods (see Methods), but use our knowledge of the simulated ground truth to produce trajectories free from false positives and linking errors which may lead to bad performance that do not reflect the intrinsic quality of the trajectory analysis. We allow at most 3 consecutive missing positions before starting an new trajectory.

As a test problem, we consider tracking of tRNA molecules in E. coli cells [22, 25], which should present several interesting difficulties. At least three discernible diffusive states may be expected: ribosome bound (slow), free (fast diffusion), and a ternary complex (intermediate). The ribosome-bound states further display spatial structure in the form of nucleoid exclusion [25], as well as non-exponential waiting times [26], since tRNA goes through several reaction steps before dissociating from the ribosome [27]. We constructed a simplified kinetic and spatial model incorporating these features and generated synthetic fluorescent microscopy data with 200 Hz frame rate [9], as shown in Fig. 4. Rate constants are chosen to give a steadystate occupancy of 20/30/50 (bound/unbound/ternary complex), and we vary the overall rates to get total bound state mean dwell time *τ*_*B*_ between 50ms and 400ms (see Methods).

**Figure 4.**
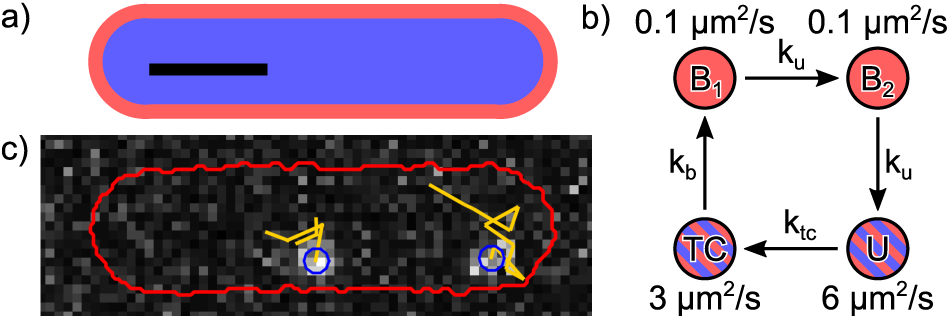
Model for simulated tracking of fluorescent tRNA molecules. (a) Cross-section of the simulation geometry, which consists of concentric cylinders with spherical end-caps, representing the nucleoid (blue) floating in the cytoplasm (red). Scale bar (black) 1 μm. (b) Kinetic model of the tRNA cycle. Two states ***B***_1_, ***B***_2_ with low diffusion coefficient represent ribosome bound states, and are excluded from the nucleoid, while the two unbound (U) and ternary complex (TC) states are free to roam the whole cell. (c) A simulated frame with two fluorophores in a single cell, with cell outline (red) and particle tracks (yellow) added. Pixels size 80 nm.

Not all estimated model parameters can be directly compared to the simulated model parameters. For example, the nucleoid exclusion means that the *TC → B* reaction cannot take place in the nucleoid region, lowering the effective value of *k*_*b*_. Nucleoid exclusion also distorts the state occupancy of the detected spot population, because defocused spots are more difficult to detect, and with our simulated focus in the cell midplane, the bound states are relatively enriched in defocused regions. In Fig. 5, we therefore focus on comparisons where these complications are minimal.

**Figure 5.**
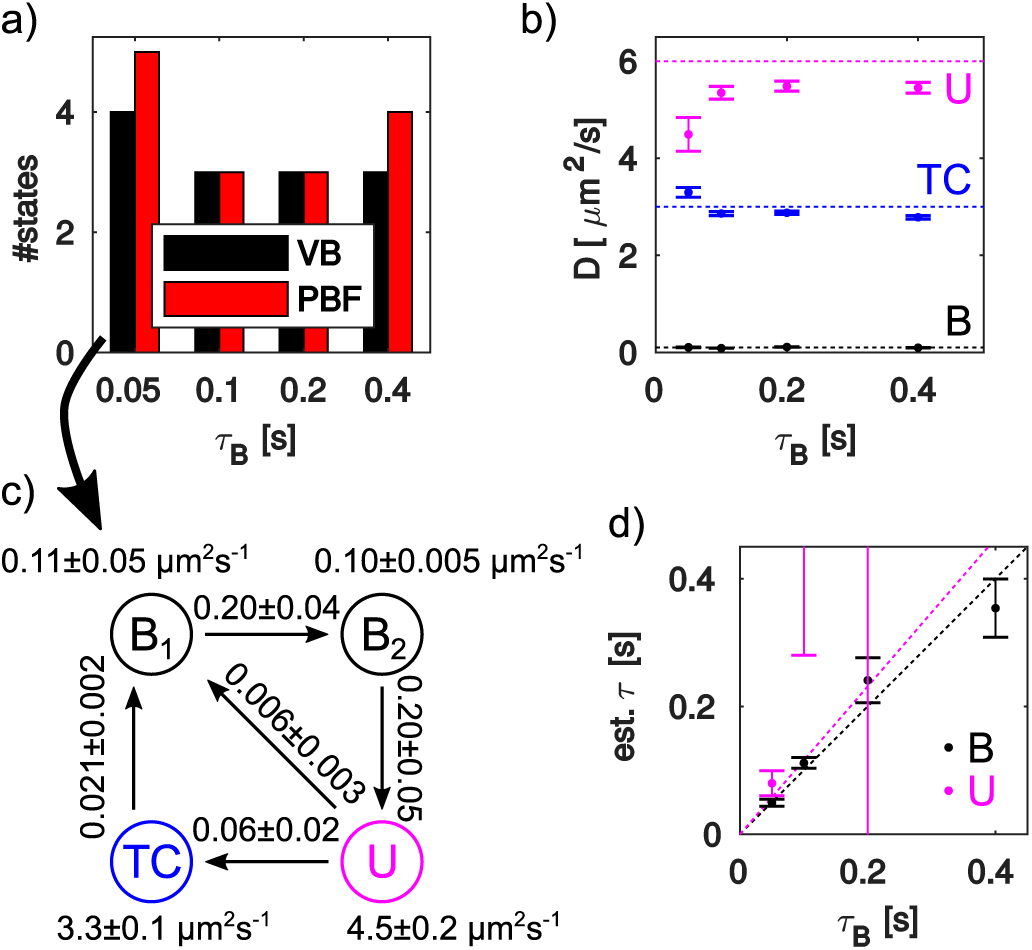
Analysis of simulated microscopy data. The different ground truth models are denoted by their total bound state dwell times, *τ*_*B*_ = 0.05s, 0.1s, 0.2s, and 0.4s, respectively. a) Number of states selected by the VB and PBF criteria. b) Diffusion coefficients for VB-selected models. Dashed colored lines indicate the true diffusion constants of the U, TC, and B1,2 states. For the 0.05 s model, two states near 0.1 μm^2^ s^−1^ are found. c) Kinetic scheme of the 4-state 0.05 s model, with transition probabilities per time-step below 10*-*^8^Δ*t*^-1^ suppressed. States are named and colored according to the obvious similarity with the true scheme in Fig. 4b. For the true model, ***k***_***u***_ = 0.2Δ*t*^-1^ and ***k***_***tc***_ = 0.067Δ*t*^-1^. d) Bound (B) and unbound (U) state mean dwell times, computed from the transition probability matrix. For the 0.05 s model, we added the dwell times of the two B states. Dashed lines indicate the true mean dwell times. All precision indicators are bootstrap standard error of the mean, and all data sets contain about 16 000 steps.

The true model contains three diffusion constants but 4 kinetic states. Starting with the number of states (Fig. 5a), we see mostly four states, and note that the VB and PBF model selection agree half the time, and that the PBF favors more states in cases of disagreement. Plotting the diffusion constants from VB-selected models (Fig. 5b), we see that it finds diffusion constants close to the true values, although for the fastest kinetics (*τ*_*B*_ = 50ms) the high-D states are biased towards each other, probably because the faster dynamics produce more short events that make it more difficult for the HMM to distinguish the two fast states correctly. There is also a general downward bias in the highest diffusion constant, perhaps because the HMM can only confuse it with slower states, or perhaps because of confinement effects.

Regarding the kinetics, a detailed look at the 4-state model for the fastest kinetics (Fig. 4c) shows a striking resemblance to the true rate model in Fig. 4b. The unidirectional cycle is clearly visible, and the transition probabilities corresponding to *k*_*u*_ and *k*_*t*_*c* closely resemble the underlying ground truth. The mean dwell times of this model is comparable to the average trajectory length of about 0.12s. For slower models, the bound state dwell time is well reproduced (5d, black), but otherwise the kinetics is not as well reproduced. Only one bound state is identified, the unbound dwell times (5d, magenta) are not close to the true values, and the transition matrices (not shown), do not resemble the cyclic pattern of the underlying model. However, even the more limited ability to measure the mean dwell time of a slow or immobile state when that dwell time exceeds the average trajectory length could be of biological interest, for example to study the interactions of small molecules such as tRNA or proteins interacting with larger structures such as ribosomes or DNA.

## II. DISCUSSION AND CONCLUSION

Together with methods to extract both positions and position uncertainty from images of single spots [7], the variational algorithm we present here makes it possible to significantly decrease analysis artifacts due to variable localization quality, such as out-of-focus motion, gradual bleaching or stage drift, or fast fluorophore blinking. We expect this to become a powerful tool for quantitative analysis of diffusive *in vivo* single particle tracking data.

Compared to our previous implementation [7], the present algorithms are significantly faster, which makes exploratory analysis of large data sets practical, and support both maximum likelihood or variational Bayes inference, which allows a more nuanced statistical analysis. Here, we compared model selection by the Akaike criterion (AIC) [21], a variational implementation of cross-validation using pseudo Bayes factors (PBF) [20], and the purely Bayesian variational Bayes (VB) approach [10, 11]. While the AIC clearly overfitted in our example, the PBF showed a much weaker preference for more complex models compared to VB, and in light of the theoretical arguments that can be made against a purely Bayesian approach in certain situations [18, 19], we think this approach merits further study.

One can also imagine combining our approach to localization uncertainty with other types of heterogeneity in the data. Interesting examples include variability in the underlying diffusion constant or other model parameters [16], or the presence complex spatial structure [28].

Another interesting direction for future work is to consider more complex models. The simple diffusion models could potentially be extended in several useful ways within the exponential family of models that enable efficient variational algorithms[10]: Localization errors couldbe treated as model parameters rather than external observations, possibly depending on the chemical state or coordinate dimension (see SI Sec. S5). Combinations of directed motion and confinement in harmonic potentials that lead to Gaussian models have been proposed for analysis of particle tracking experiments [3, 29]. Introducing explicit termination rates could correct bias that arise from correlations between chemical states and trajectory termination [30], for example when fast diffusing molecules move out of focus faster than slow diffusing ones [31].

### Software

Our algorithms are freely available as open source Matlab code from https://github.com/bmelinden/vbSPTu. The vbSPTu software suite includes a GUI to run a simple standard analysis, support for scripting large analysis tasks, and low-level tools for creating customized analysis.

## METHODS

### B Simulated trajectories

For the model selection experiments in Figures 1 and Figure S1, we used synthetic trajectories simulated using the analysis model, Eqs. (2-4).

We simulated a 3 state model with the following parameters: Diffusion constants *D*_1_ = 0.1μm^2^ s^−1^, *D*_2_ = 6μm^2^ s^−1^, and *D*_3_ = 3μm2 s^−1^. The kinetic model is an irreversible cycle *D*_1_ → *D*_2_ → *D*_3_ → *D*_1_ → … with mean waiting times of 100 ms, 100 ms, and 116 ms, respectively. The positions are simulated according to Eqs. (2 -4), with motion blur corresponding to an exposure time of *t*_*E*_ = 1.5ms (*τ* = 0.15, *β* = 0.0775).

For the static localization errors *v*_*t*_, we used a phenomenological model of defocus-widening. First, we confined 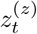 (the *z* component of *z*_*t*_) to 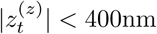 nm using the method of images. Then, we defined an effective spot width

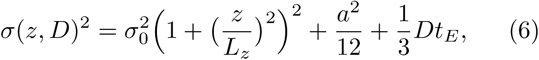

with *σ*_0_ = 135nm, *L*_*z*_ = 300nm, *a* = 80nm. The *a*^2^ term approximates the effect of finite pixel size [32], and the *Dt*_*E*_ term spotwidening due to motion blur [6]. We then computed *v*_*t*_ using the approximate Cramer-Rao lower bound expression of Mortensen et al. [32],

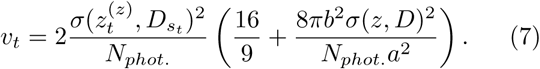

for use in Eq. (4). We analyzed the *x* and *y* components of *x*_*t*_, and chose trajectories to be exponentially distributed with mean length 25, but discarded trajectories with length below 5. For Fig. 3a,b, we used a data set of 100 trajectories. For the statistical model selection study (Figure 1 and Figure S1), we sampled data sets of various sizes from a large data set of several hundred thousand positions, such that all model selection techniques used the same set of trajectories.

### C. Simulated microscopy

Simulated video-microscopy images for tRNA tracking was generated using the SMeagol simulation software [9] with the spatial reaction diffusion model illustrated in Fig. 4. We simulated uniform exposure during 1.5 ms of the 5 ms sampling time. Camera noise was generated using a high gain approximation of EMCCD noise [32] with offset 200, EM gain 77, and Gaussian readout noise with standard deviation 20. We used 80 nm pixels and a uniform background fluorescence that decayed from two to one photon/pixel with a decay rate of 2 s^−1^. For the optics, we used a Gibson Lanni pointspread function (PSF) model [33] generated by PSFgenerator [34], with *λ* = 680nm and *NA* = 1.49.This is a spherically symmetric PSF, suitable for isotropic emitters or fluorophores with high rotational mobility. Fluorescent spot intensity was set to give on average of 200 photons per frame, and the average bleaching time was 20 frames. Using custom Matlab scripts, we simulates 200 frame movies with about 30 cells spread evenly across a 512 × 130 pixel field of view, with a few active fluorescent spots per cell. An example of one such cell is shown in Fig. 4c.

### D. Spot detection and localization

We use the fast radial symmetry transform[35] for spot detection, and estimated spot positions and localization uncertainty using a symmetric Gaussian spot model and maximum aposteriori estimates on 9by9 regions of interests, as described by Lindén et al. [9].Spots with 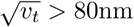 80nm were discarded from the analysis.

## Acknowledgements

This project was funded by grants to J.E. from the Knut and Alice Wallenberg Foundation and the European Research Council.

## Author contributions

M.L. and J.E. planned research and wrote the paper. M.L. designed and implemented new analysis methods, generated data, and analyzed data.

